# A “Cell-Nucleus Segmentation” Script for Non-Invasive Nuclear Dry Mass Measurement

**DOI:** 10.1101/2020.10.28.358382

**Authors:** Edward M. Kong, Chenfei Hu, Byoung Soo Kim, Gabriel Popescu

## Abstract

The nucleus is the largest organelle in cells carrying genetic materials that support genetic replication and transcription. It is likely that such genetic activities can influence the nuclear *dry mass*, but there is a lack of analytical tools enabling us to monitor dynamic changes in this quantity. To this end, this study demonstrates an image analysis script that allows us to quantify these changes in the nuclear dry mass. The script runs the cell-nuclei segmentation using Matlab. By using the fluorescent image as a template for the boundaries of cell nuclei and quantitative phase images for retrieving the dry mass density, the script recognizes nuclei of all cells in an image at a time and quantifies the nuclear dry mass. Using the “the cell-nucleus segmentation” script, this study reveals an interesting correlation between the nuclear dry mass and the filopodia protrusion of cervical epithelial cells. As the filopodia density and protrusion length increase, the nuclear dry mass increases. On the other hand, whenever the nuclear dry mass decreases, cells filopodia retract significantly. Taken together, the imaging script developed here will be useful to quantifying dynamic nuclear activities of a broad array of cells non-invasively.

## Introduction

The cell nucleus is enclosed by self-associated phospholipid bilayers. According to previous studies, the cell nucleus undergoes significant shape and size changes during various phenotypic events. For example, cell polarization and migration result in complex nuclear shape change from ellipsoidal to cigar-like shape, depending on cytoskeletal dynamics, surrounding tissue porosity, and migration mode. Differentiation of mesenchymal stem cells to adipocytes also decrease the nuclear area significantly while making the nucleus more elongate (1). These shape changes are caused by membrane protein synthesis or force action on the nuclear envelope (2–4). Thus, cell nucleus shape becomes a marker of various injuries and diseases because depletion of genetic materials in the nucleus alters interaction between nuclear envelope and interior cytoskeletons and, in turn, deforms the nucleus (5).

In addition, it has been suggested that such cellular nuclear shape changes would result from changes of the non-aqueous content in the nuclear, referred to as dry mass (6). The dry mass represents the amount of genetic material in the cell nucleus (7). Such change in the dry mass may further influence the nuclear shape. However, this insight has been remained hypothetical, likely due to the lack of analytical tools enabling us to monitor dynamic changes in the dry mass. Our group has developed a quantitative phase imaging (QPI) techniques that combine phase contrast microscopy with phase-shifting interferometry to track total cell dry mass change (8–13). Cell proliferation tends to increase the dry mass, while apoptosis induces the opposite effect. Such analysis can be extended to assess nuclear dry mass, but a method enabling us to quantify it non-invasively is lacking.

To this end, this study demonstrates an imaging analysis script that allows us to quantify the nuclear dry mass from the quantitative phase image of cells. The script was made to run the cell-nuclei segmentation using Matlab. By using the fluorescent image as a template for the boundaries of cell nuclei and phase images for retrieving the dry mass density, the script recognizes the nuclei of all cells in the field of view (QPI). The script sums the pixel phase values of each cell to quantify the dry mass. This script can continue to analyze the nuclear dry mass of cells in a multiple number of images. In this study, we use the cell-nuclei segmentation script to quantify the nuclear dry mass of HeLa cells, a cervical epithelial cell line. We further examined if the nuclear dry mass is related to the filopodia protrusion activities (14, 15). Overall, this study serves to provide a useful method to monitor dynamics of various cell nuclear events in a non-invasive and real-time manner.

## Methods

### Quantitative phase imaging of HeLa cells

HeLa cells were seeded onto a glass-bottom dish and cultured in high glucose DMEM supplemented with 10 % fetal bovine serum, 100 U/ml penicillin, and 0.2 mg/ml streptomycin (all from GIBCO-BRL Life Technologies, Grand Island, NY) under standard conditions at 37°C, 5% CO_2_. The nuclei of cells were stained with (4’,6-diamidino-2-phenylindole) (DAPI) (1:1,000 dilution), and the culture media were exchanged to CO_2_ independent medium (18045-088, Thermo Fisher). Then, the cells were imaged using spatial light interference microscopy (SLIM)(16). DAPI-stained nuclear fluorescent image and quantitative phase maps were acquired. Nuclear dry mass of individual cells captured in the images was analyzed using the cell-nuclei segmentation script that will be explained in the result section.

## Results and Discussion

### Script developed to monitor nuclear dry mass

We examined if the cell-nuclei segmentation script can analyze changes in the nuclear dry mass quantitatively. The principle of segmentation method is summarized in Fig. 1. First, nuclei stained with the UV excitable Hoechst 33342 and quantitative phase images of cells were captured. Then, the first fluorescent image of the cell nuclei (Fig. 2a) is converted to a grayscale image using the Matlab syntax “rgb2gray” (Fig. 1 and Fig. 2b). The RGB images contain R, G, and B components, each of which contain information about the individual gradient colors. With R, G, and B components, we could take a weighted sum and use it for color-to-grayscale conversion.

**Figure 1.**
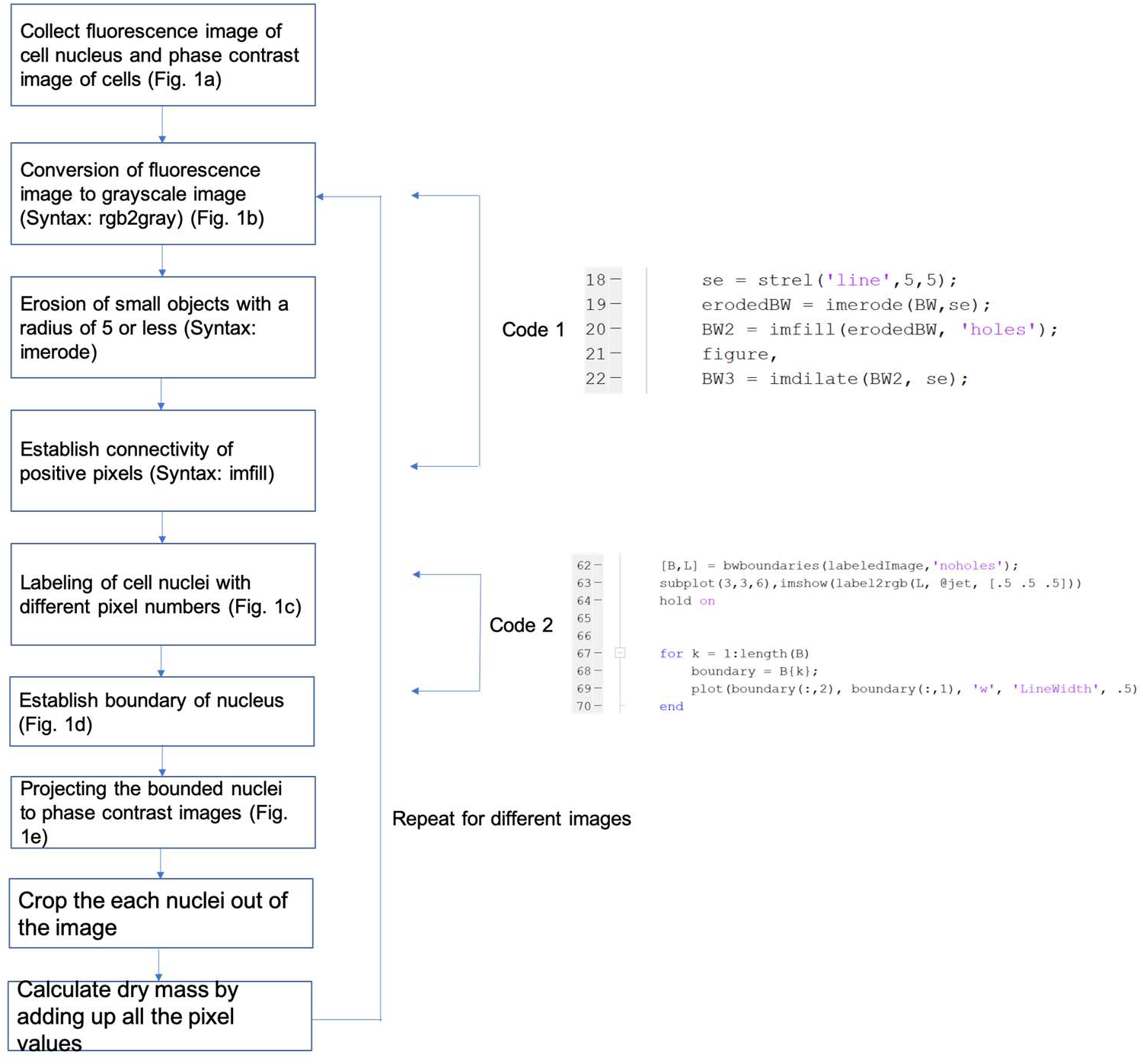
Schematic description of the cell-nuclei segmentation script used to measure nuclear dry mass.

**Figure 2.**
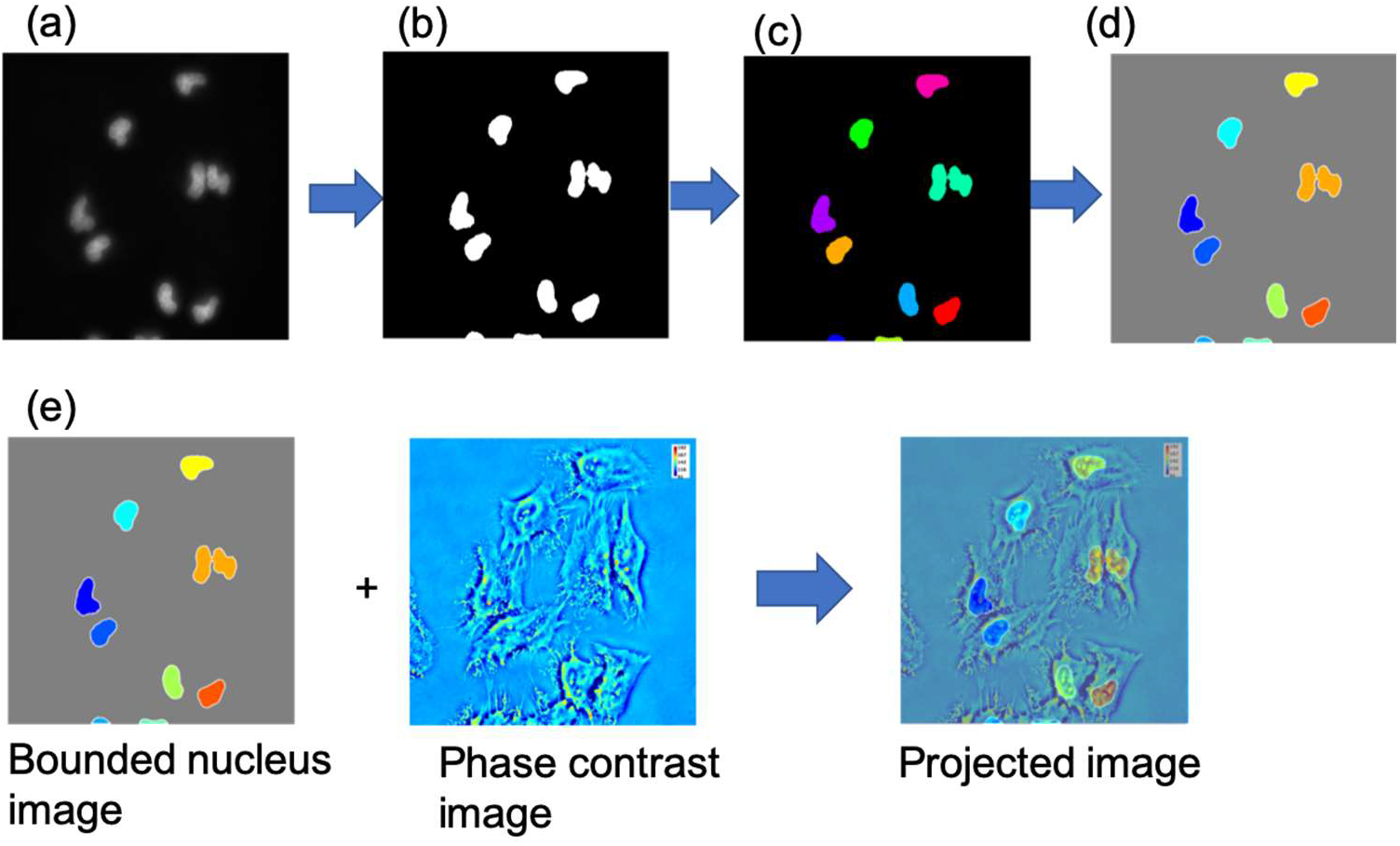
Cell nuclei and quantitative phase images processed by the cell-nuclei-segmentation script. (a) Fluorescence image of cell nuclei, (b) grayscale image, (c) labeled image, (d) bounded image, and (e) projection of the bounded nuclei to the phase contrast image of cell.

Next, using the built-in Matlab syntax “imbinarize”, the script binarizes the greyscale images by converting every pixel value to 0 or 1 (Fig. 2b). The pixel values are rounded to 1 if they are positive and rounded to 0 if they are nonpositive. The script also includes “imfill” that tracks connectivity of pixels and fills in “zero” pixel values with “one” pixel values, to increase connectivity and fill in the holes. Next, the script counts connectivity of each pixel cluster (i.e., pixels touching each other and diagonal to each other). If connectivity is less than 50 pixels, then the cluster is converted to all zero-value pixels. The BW label syntax once again counts the connectivity and counts each separate body as morphological “objects”. It outputs a matrix that labels the cells with different pixel numbers as shown in Fig. 2c with color.

Finally, the script determines a matrix containing the boundary pixel coordinates, marked with “B”, using the syntax. “bwboundaries” This process is repeated to establish a boundary for each cell. Matlab uses a contour tracing algorithm called the Moore-Neighbor tracing algorithm. Denote *f*(*x*) the set of 8 pixels that surround the input pixel “*x*” (also known as Moore neighborhood). The algorithm moves from bottom to top and left to right until the first positively valued pixel is found. This is put into the set B (where B contains all the pixels representing a boundary). The algorithm then finds the next clockwise pixel within the Moore neighborhood, and declares that pixel as the next boundary pixel, as well as the new “*x*”. This loop will end when the final clockwise pixel detected is the first pixel that was entered initially. The syntax “bwboundaries” repeats this loop for each connected body in the image. The script then uses these coordinates and label matrices (from “bwboundaries”) to establish a bounding box (Fig. 2d).

These bounding boxes/coordinate descriptions and locations are projected onto the phase image. Each cell nuclei is then cropped out of the image and used to calculate dry mass by adding up all the pixel values in the specific phase contrast image (Fig. 2e). We considered the sum of the pixel values within the outline as the dry mass coefficient. The loop calculates each cell’s dry mass on the phase image. The script then loops back to the next set of images consisting of one phase contrast image at each time point.

### Analysis of nuclear dry mass of cells with different filopodia protrusion activities

We first captured quantitative phase images (QPI) of the HeLa cells cultured in a growth media. As shown in Fig. 3, cells protrude multiple number of filopodia, which are formed by tightly bundled actin filaments. The number and protrusion length of filopodia quantified using the ImageJ software increased over time. There were no significant changes of the cell spreading area and nuclear shape was made during the same period. Interestingly, during the same period, average nuclear dry mass quantified with cell nuclei isolated from the QPI images using the cell-nuclei isolation script was increased by more than two-fold (Fig. 4a). Overall, the nuclei dry mass was related to the number and protrusion length of filopodia (Fig. 4b).

**Figure 3.**
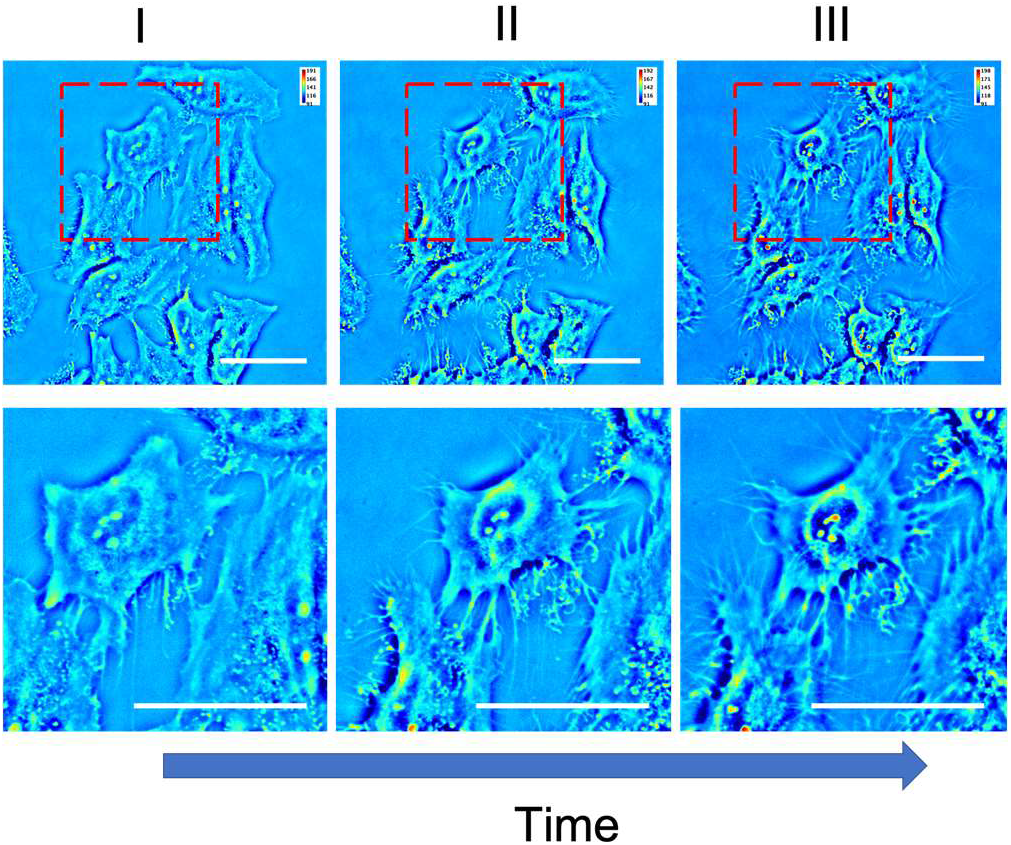
Quantitative phase images of HeLa cells. Images (I-III) show increases of filopodia density and protrusion of cells over time. Images on the 2^nd^ row are magnification of the area marked with red dotted box. Scale bars represent 10 μm.

**Figure 4.**
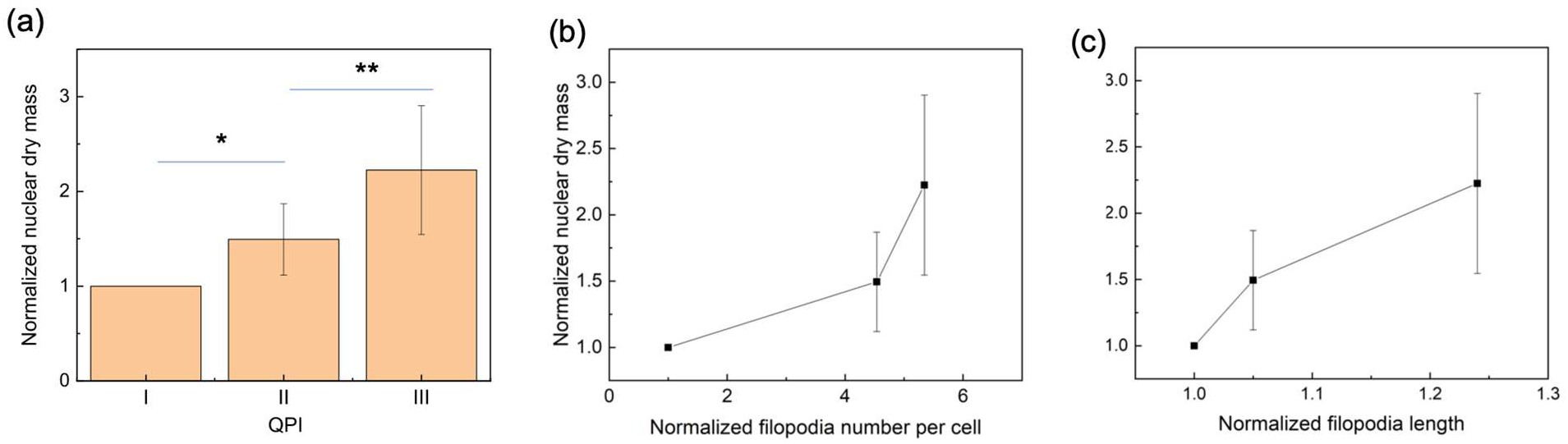
(a) Nuclear dry mass of individual cells quantified by processing the images in Fig. 2 using the cell-nuclei segmentation script. The nuclear dry mass at each stage was normalized to the nuclear dry mass of cells shown in the Fig. 2I. Values and error bars represent the average and standard deviation of 10 different cells per image. * and ** represent statistical significance of the differences between each value (*p < 0.05 & **p < 0.05). (b) Correlation of normalized nuclear dry mass to the normalized filopodia number per cell. The filopodia number was normalized to the filopodia number of cells in Fig. 2I. (c) Correlation of normalized nuclear dry mass to the normalized filopodia protrusion length. The average protrusion length was normalized to the protrusion length of filopodia shown in Fig. 2I.

Next, we examined if the filopodia retraction is also correlated to the nuclear dry mass. According to the quantitative phase contrast image of cells shown in Fig. 4a, cells undertake significant decrease in the number of protruded filopodia by 15 folds. Accordingly, the average nuclear dry mass quantified using the cell-nuclei segmentation script decreased by 1.7-fold (Fig. 5b). Therefore, the decrease of nuclear dry mass was correlated to the number of filopodia per cell (Fig. 5c).

**Figure 5.**
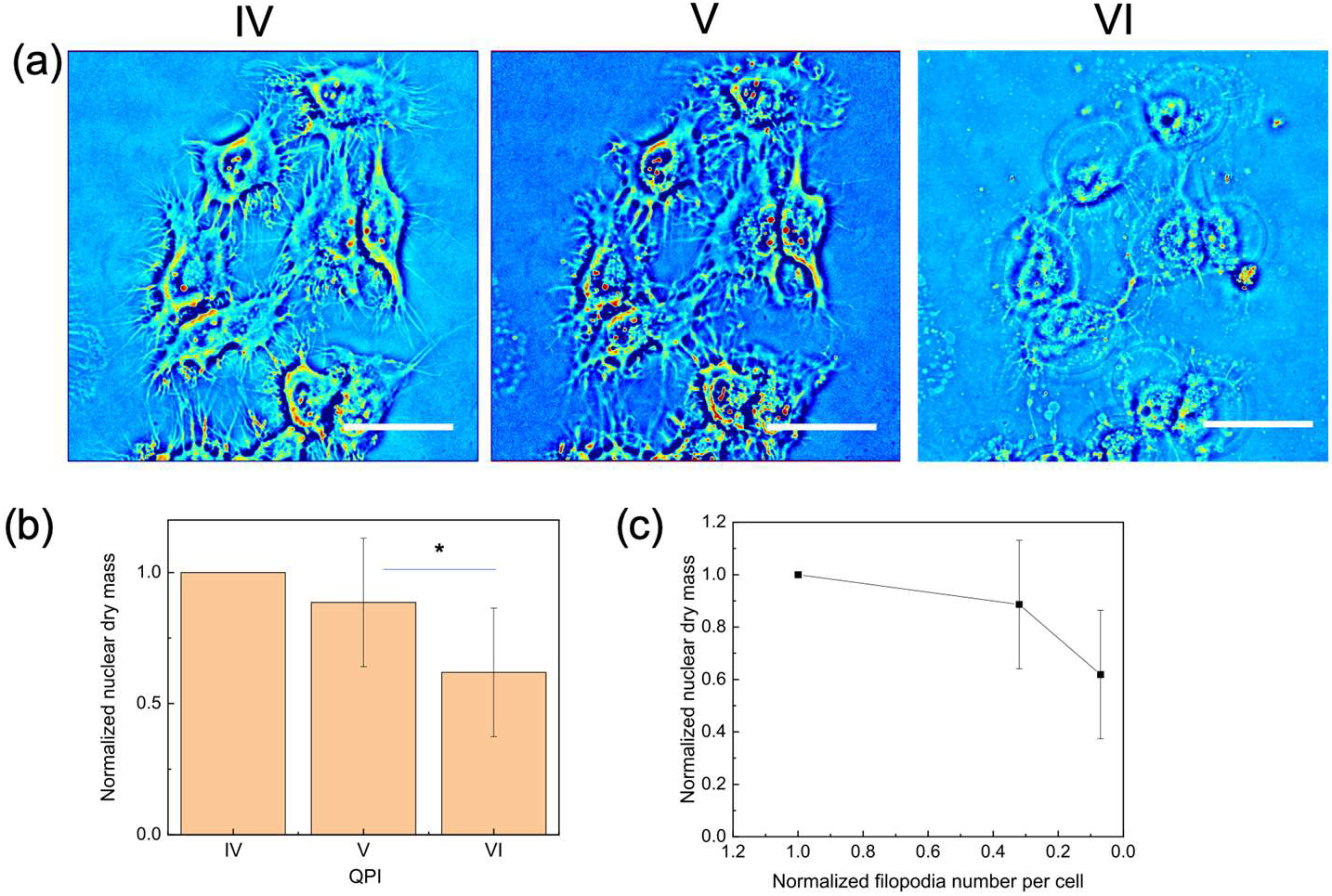
(a) Quantitative phase images of HeLa cells. Images (IV-VI) show decrease of filopodia density over time. Images Scale bars represent 10 μm. (b) Nuclear dry mass of individual cells quantified by processing the images in (a) using the cell-nuclei segmentation script. The nuclear dry mass at each stage was normalized to the nuclear dry mass of cells shown in the (a)-IV. Values and error bars represent the average and standard deviation of 10 different cells per image. * represents statistical significance of the differences between each value (*p < 0.05). (c) Correlation of normalized nuclear dry mass to the normalized filopodia number per cell. The filopodia number was normalized to the filopodia number of cells in (a)-IV.

These results confirm that the cell-nucleus segmentation script allows us to monitor nuclear dry mass of cells captured with the quantitative phase contrast microscopy. More interestingly, the resulting changes in the nuclear dry mass were correlated to cellular activities to protrude and retract filopodia. This result suggests that actin filaments bundled to cause the membrane protrusion increase tension of cytoplasmic and perinuclear actin cables (17). These stably interconnected actin networks likely provide mechanical support for nuclear activities including replication and transcription (18). As a consequence, cells can increase mass of genetic materials within their nuclei. Conversely, retraction of filopodia would decrease tension of perinuclear actin cables. Thus, cell nucleus surrounded by softened actin networks becomes inactive for replication and transcription. It is also likely that nucleus release mRNA molecules during this stage. We propose that such nuclear dry mass analysis can be useful to analyzing nuclear activities of a broad array of cells in a non-invasive manner.

## Summary and Discussion

In sum, the results of this study demonstrate that the “cell-nucleus segmentation” script is useful in analyzing the nuclear dry mass of cells in a non-invasive manner. The script could determine the dry mass by establishing the nuclear boundary marked with fluorophores and projecting them onto the phase contrast image of the cell automatically. Thus, the script allowed us to quantify the nuclear dry mass of individual cells automatically. Using the script, we could demonstrate that dynamic filopodia protrusion and retraction influence nuclear dry mass of cells, likely due to the changes in the tension of perinuclear actin networks and subsequent nuclear positioning. We envision that the “cell-nucleus segmentation” script would be useful to studying dynamic nuclear activities of a broad array of biological cells.

## Notes

### Competing Interest Statement

The authors have declared no competing interest.

## References

1. McColloch A, Rabiei M, Rabbani P, Bowling A, Cho M. Correlation between Nuclear Morphology and Adipogenic Differentiation: Application of a Combined Experimental and Computational Modeling Approach. Sci Rep. 2019;9(1):16381. Epub 2019/11/11. doi: 10.1038/s41598-019-52926-8. PubMed PMID: 31705037; PMCID: PMC6842088.

2. Walters AD, Bommakanti A, Cohen-Fix O. Shaping the nucleus: factors and forces. J Cell Biochem. 2012;113(9):2813–21. Epub 2012/05/09. doi: 10.1002/jcb.24178. PubMed PMID: 22566057; PMCID: PMC3471212.

3. Friedl P, Wolf K, Lammerding J. Nuclear mechanics during cell migration. Curr Opin Cell Biol. 2011;23(1):55–64. Epub 2010/11/27. doi: 10.1016/j.ceb.2010.10.015. PubMed PMID: 21109415; PMCID: PMC3073574.

4. Webster M, Witkin KL, Cohen-Fix O. Sizing up the nucleus: nuclear shape, size and nuclear-envelope assembly. J Cell Sci. 2009;122(Pt 10):1477–86. Epub 2009/05/08. doi: 10.1242/jcs.037333. PubMed PMID: 19420234; PMCID: PMC2680097.

5. Dasso M, Fontoura BMA. Editorial overview: The cell nucleus: Dynamic interplay of shape and function. Curr Opin Cell Biol. 2018;52:iv–vi. Epub 2018/06/06. doi: 10.1016/j.ceb.2018.05.005. PubMed PMID: 29866401; PMCID: PMC6347106.

6. Damania D, Subramanian H, Backman V, Anderson EC, Wong MH, McCarty OJ, Phillips KG. Network signatures of nuclear and cytoplasmic density alterations in a model of pre and postmetastatic colorectal cancer. J Biomed Opt. 2014;19(1):16016. Epub 2014/01/21. doi: 10.1117/1.Jbo.19.1.016016. PubMed PMID: 24441943; PMCID: PMC4019418.

7. Davies HG, Deeley EM. An integrator for measuring the dry mass of cells and isolated components. Exp Cell Res. 1956;11(1):169–85. Epub 1956/08/01. doi: 10.1016/0014-4827(56)90201-4. PubMed PMID: 13356838.

8. Mir M, Bergamaschi A, Katzenellenbogen BS, Popescu G. Highly sensitive quantitative imaging for monitoring single cancer cell growth kinetics and drug response. PLoS One. 2014;9(2):e89000. Epub 2014/02/22. doi: 10.1371/journal.pone.0089000. PubMed PMID: 24558461; PMCID: PMC3928317 following conflicts: G. Popescu has financial interests in Phi Optics, Inc., a company developing quantitative phase microscopes, which, however, did not sponsor this research. This does not alter the authors’ adherence to all the PLOS ONE policies on sharing data and materials.

9. Popescu G, Park Y, Lue N, Best-Popescu C, Deflores L, Dasari RR, Feld MS, Badizadegan K. Optical imaging of cell mass and growth dynamics. Am J Physiol Cell Physiol. 2008;295(2):C538–44. Epub 2008/06/20. doi: 10.1152/ajpcell.00121.2008. PubMed PMID: 18562484; PMCID: PMC2518415.

10. Hu C, Popescu G. Quantitative Phase Imaging: Principles and Applications. Label-Free Super-Resolution Microscopy: Springer; 2019. p. 1–24.

11. Hu C, Popescu G. Quantitative Phase Imaging (QPI) in Neuroscience. IEEE Journal of Selected Topics in Quantum Electronics. 2019;25(1):1–9. doi: 10.1109/JSTQE.2018.2869613.

12. Popescu G. Quantitative phase imaging of cells and tissues. New York: McGraw-Hill; 2011. xx, 362 p. p.

13. Hu C, Sam R, Shan M, Nastasa V, Wang M, Kim T, Gillette M, Sengupta P, Popescu G. Optical excitation and detection of neuronal activity. J Biophotonics. 2018:e201800269. Epub 2018/10/13. doi: 10.1002/jbio.201800269. PubMed PMID: 30311744.

14. Jacquemet G, Hamidi H, Ivaska J. Filopodia in cell adhesion, 3D migration and cancer cell invasion. Curr Opin Cell Biol. 2015;36:23–31. Epub 2015/07/18. doi: 10.1016/j.ceb.2015.06.007. PubMed PMID: 26186729.

15. Mattila PK, Lappalainen P. Filopodia: molecular architecture and cellular functions. Nat Rev Mol Cell Biol. 2008;9(6):446–54. Epub 2008/05/10. doi: 10.1038/nrm2406. PubMed PMID: 18464790.

16. Wang Z, Millet L, Mir M, Ding H, Unarunotai S, Rogers J, Gillette MU, Popescu G. Spatial light interference microscopy (SLIM). Opt Express. 2011;19(2):1016–26. Epub 2011/01/26. doi: 10.1364/OE.19.001016. PubMed PMID: 21263640; PMCID: PMC3482902.

17. Huelsmann S, Brown NH. Nuclear positioning by actin cables and perinuclear actin: Special and general? Nucleus. 2014;5(3):219–23. Epub 2014/06/07. doi: 10.4161/nucl.29405. PubMed PMID: 24905988; PMCID: PMC4133217.

18. Kim DH, Wirtz D. Cytoskeletal tension induces the polarized architecture of the nucleus. Biomaterials. 2015;48:161–72. Epub 2015/02/24. doi: 10.1016/j.biomaterials.2015.01.023. PubMed PMID: 25701041; PMCID: PMC4917004.

